# Stomatal responses at different vegetative stages of selected maize varieties of Bangladesh under water deficit condition

**DOI:** 10.1101/2021.09.13.460018

**Authors:** Md. Moin Uddin Talukder, Pinky Debnath, Sonia Nasrin, Sonia Akter, Md. Raihan Ali, Md. Rejaul Islam, S M Abdul-Awal

## Abstract

Drought stress causes stomatal behavior change in most plants. Water deficit condition caused by drought is one of the most significant abiotic factors reducing plant growth, development, reproductive efficiency, and photosynthesis, resulting in yield loss. Maize (Zea mays L.) holds a superior position among all the cereals due to its versatile use in the food, feed, and alcohol industries. A common demonstrative feature of a complex network of signaling pathways led by predominantly abscisic acid under drought conditions is stomatal aperture reduction or stomatal closure, which allows the plant to reduce water loss through the stomatal pore and to sustain a long time on water deficit condition. This study analyses the stomatal density, stomatal closure percentages, and guard cell aperture reduction using a microscopy-based rapid & simple method to compare guard cell response & morphological variations of three hybrid maize varieties viz. BHM (BARI hybrid maize)-7, BHM-9, and BHM-13 developed by Bangladesh Agricultural Research Institute (BARI). A drought treatment was applied to all varieties at two different vegetative stages, vegetative stage 3 (V3) and V5, until they reach V4 and V6, respectively. After drought exposure at the V4 stage, the percentage of closed stomata of BHM-7, BHM-9, and BHM-13 was 21%, 23%, and 33%, respectively. The reduction in the guard cell aperture ratio of BHM-7, BHM-9, and BHM-13 was 14.83%, 10.92%, and 33.85%, respectively. At the V6 stage, for the second set of plants, the closed stomata of BHM-7, BHM-9, and BHM-13 were 18%, 21%, and 34%, respectively. The rate of reduction in guard cell aperture ratio of BHM-7, BHM-9, and BHM-13 was 5.52%, 2.48%, and 38.75%, respectively. Therefore, BHM-13 showed maximum drought adaptation capacity compared to BHM-7 and BHM-9 due to the highest percentage of closed stomata and the highest percentage of reduction in aperture ratio.

## Introduction

Globally, drought is the most harmful environmental phenomenon that comes with financial hardship among farmers in developed countries, malnutrition, and even famine in third-world countries (1). It adversely affects almost every physiological process in the plant, such as membrane fluidity and function, decreases photosynthesis, causes injury, aberrant physiology, limitation of growth, and increases susceptibility to insects and disease-causing pests (2). Plants utilize various defense mechanisms to endure drought stress. Plants usually close their stomata in response to the environment. For instance, most plants close stomata at night, while plants may also close their stomata under severe conditions such as drought to limit the amount of water evaporation from their leaves. The opening and closing of stomata are a very well-regulated masterpiece of plant evolution driven by the translation of chemical signals into the mechanical movement of guard cells. Stomata are formed by two specialized guard cells, morphologically distinct from general epidermal cells (3). Guard cells surround stomata pores in the epidermis of plant leaves and stems. Stomata allow the diffusion of CO2 into the leaf for photosynthesis and the diffusion of H2O out of the leaf during transpiration (4). Plants lose over 95% of their water content via transpiration to the atmosphere. Pairs of guard cells regulate this gaseous exchange.

During water deficit conditions, water loss through transpiration is reduced in response to abscisic acid by promoting stomatal closure and inhibition of opening (5). Various environmental factors such as drought, CO2 concentration, light, humidity, biotic stresses, and different plant hormones regulate stomatal apertures (6). These changes are driven by cation and anion effluxes. Opening or closure of stomata is achieved by osmotic swelling or shrinking of guard cells, respectively, driven by transmembrane ion fluxes of K+, Cl-, and malate2- (7). Reorganization of the cytoskeleton, metabolite production, posttranslational modifications of existing cellular proteins, and modulation of gene expression are critical components of guard cell biology and determinants of stomatal regulation (8). Various major hormones are involved in stomatal regulation. Among these, ABA plays the overriding role (9). Increased level of ABA concentrations induces multiple cascades of biochemical events like protein phosphorylation, generation of nitric oxide (NO) and hydrogen peroxide (H2O2), changes in intracellular Ca2+ concentration, and membrane depolarization (10), leading to the modifications in the activity of ion channels, decrease of the osmotic pressure in guard cells and, thereby, closure of stomata (6).

Other phytohormones, namely ethylene, jasmonates, and salicylic acid, also function in modulating stomatal aperture. Signaling pathways triggered by hormones and pathogen attacks often involve the generation of second messengers like NO and H2O2 (11). H2O2 has a marked effect on stomatal aperture. H2O2 can induce stomatal closure and inhibit stomatal opening without any damage to cells (12). Histidine kinase AHK5 is involved in one pathway by which H2O2, derived from endogenous and environmental stimuli, is sensed and transduced to affect stomatal closure (13). Arabidopsis mutants lacking AHK5 show reduced stomatal closure in response to H2O2, which is reversed by complementation with the wild-type gene. Auxin regulates stomatal opening positively, although it can also inhibit stomatal opening when applied exogenously at high concentrations. Auxin promotes the activation of plasma membrane-localized H+-ATPases, leading to a hyperpolarization of the membrane. Activation of the K+-channels mediates an influx of potassium ions, followed by the stomata’s opening. Here, we apply a microscopy-based technique to compare the percentage of close stomata and reduction in aperture ratio among the three maize varieties under water deficit conditions. BHM-13 showed the highest percentage of closed stomata and reduced aperture ratio compared to the other two varieties (BHM-7 and BHM-9), indicating that BHM-13 has a high water utilization capacity during drought condition by reducing excess water loss.

## Materials & Methods

### Seed germination and plant growing

Seeds of three different varieties (BHM-7, BHM-9, and BHM-13) were transferred to the Petri dish containing blotting paper. The Petri dishes were placed in the growth chamber (Temperature: 25 ± 1°C, Humidity: 65%, Light: 2000 lux, Photoperiod: 12-hour light: 12-hour dark) for five days. Water was sprayed on the petri dish once a day for germination. Germinated seeds of each variety were transferred into the plastic bag containing soil and fertilizers. Soil was prepared & mixed with fertilizers like TSP, MOP, Gypsum, according to Hand Book of Agricultural Technology by Bangladesh Agricultural Research Council (14). An equal amount of soil-containing fertilizers was taken into the plastic bag. Germinated seeds were transferred into each plastic bag.

### Drought experiment

After sowing germinated seeds into the plastic bag, regular watering was provided to each bag manually once a day to allow the seeds to grow until vegetative stage 3 (V3) and vegetative stage 5 (V5). Water was sprayed once a day to half of the plants from both stages (V3 and V5; control plants), and the other half of the plants from both stages (V3 and V5) were kept without water (treated plants) until they reached V4 and V6 stage, respectively. After drought exposure, both control and treated plants received water until they reached at V5 and V7 stages.

### Isolation of epidermal peel

A simple, cost-effective method was developed to isolate epidermal peel without using safranin and glycerol solution. A scissor was used to slightly cut the plant’s abaxial surface of the second leaf of the plant. Sharp forceps were used to take off a small portion of the leaf epidermal peel (<0.5 cm) as stomata are located in this area. The leaf epidermis was placed in a glass slide and a drop of water was added to the epidermis. The glass slide was then covered by a coverslip. Epidermal peel was collected at V3 and V5 stage just before drought exposure, at V4 and V6 stage just after drought exposure, and at V5 and V7 stage after re-watering of both control and treated plants.

### Stomatal density measurement

Stomata were detected in the epidermal peel using Carl Zeiss Microscope affiliated with Zen Blue 2.0 software and counted per mm^2^ by ImageJ software. Stomata of three leaves of each variety were chosen randomly. Stomatal closure was calculated, and the final stomata open and closure percentage were determined with respect to the total stomata number per mm2.

### Stomatal closure and open percentage measurement

A specific area was measured by ImageJ software by using the scaling of the Zen Blue 2.0 software. Then the number of close and open stomata was counted on that area. After that, the probable total number of closed and open stomata was estimated mathematically in mm2 of leaf area.

### Measurement of guard cell aperture

The guard cell aperture ratio was counted by the width/length method described by Russell Johnson (15). Open stomata will have width/length values of 0.29 or more, partially open stomata will have 0.18 to less than 0.29, and fully closed stomata will have 0. Specific numbers of stomata were taken randomly per mm2 to measure the guard cell aperture ratio in the second leaf of plants in each variety at different vegetative stages.

### Revival capacity measurement

The stomatal ratio was measured for the control (regularly watered) and treated group (drought treated followed by re-watering). Then a comparison was conducted to deduce which variety has the lowest aperture ratio difference between the control and treated groups. Variety with the lowest aperture ratio difference has the best revival capacity because they are close to the standard plant (control group).

### Statistical Analysis

Data were analyzed using GraphPad Prism 8.0 software. The student’s t-test was used to compare different stages of two groups & One-way ANOVA with Tukey-Kramer post-test was used for multiple comparisons.

## Results

### Stomatal density among the varieties

Before drought stage at V3 stage, BHM-7, BHM-9, and BHM-13 had a stomatal number of 206 ± 2.18 (mean ± SE), 198 ± 2.02 (mean ± SE), and 208 ± 1.15 (mean ± SE), respectively (Fig. 1A) whereas, at V5 stage, BHM-7, BHM-9, and BHM-13 had a stomatal number of 334 ± 1.73 (mean ± SE), 319 ± 8.02 (mean ± SE), and 345 ± 6.57 (mean ± SE), respectively (Fig. 1B). After drought exposure at the V4 stage, the stomatal number of BHM-7, BHM-9, and BHM-13 was 336 ± 4.163 (mean ± SE), 308 ± 6.009 (mean ± SE), and 340 ± 4.163 (mean ± SE), respectively (Fig. 1C) whereas, at V6 stage, the stomatal number of BHM-7, BHM-9, and BHM-13 was 357 ± 3.756 (mean ± SE), 314 ± 2.728 (mean ± SE), 321 ± 13.58 (mean ± SE), respectively (Fig. 1D). After re-watering at the V5 stage, the stomatal number of BHM-7, BHM-9, and BHM-13 was 362 ± 18.78 (mean ± SE), 364 ± 67.59 (mean ± SE), and 338 ± 24.79 (mean ± SE), respectively (Fig. 1E) whereas, at V7 stage, the stomatal number of BHM-7, BHM-9, and BHM-13 was 278 ± 5.364 (mean ± SE), 326 ± 3.055 (mean ± SE), and 356 ± 2.963 (mean ± SE), respectively (Fig. 1F).

**Fig 1.**
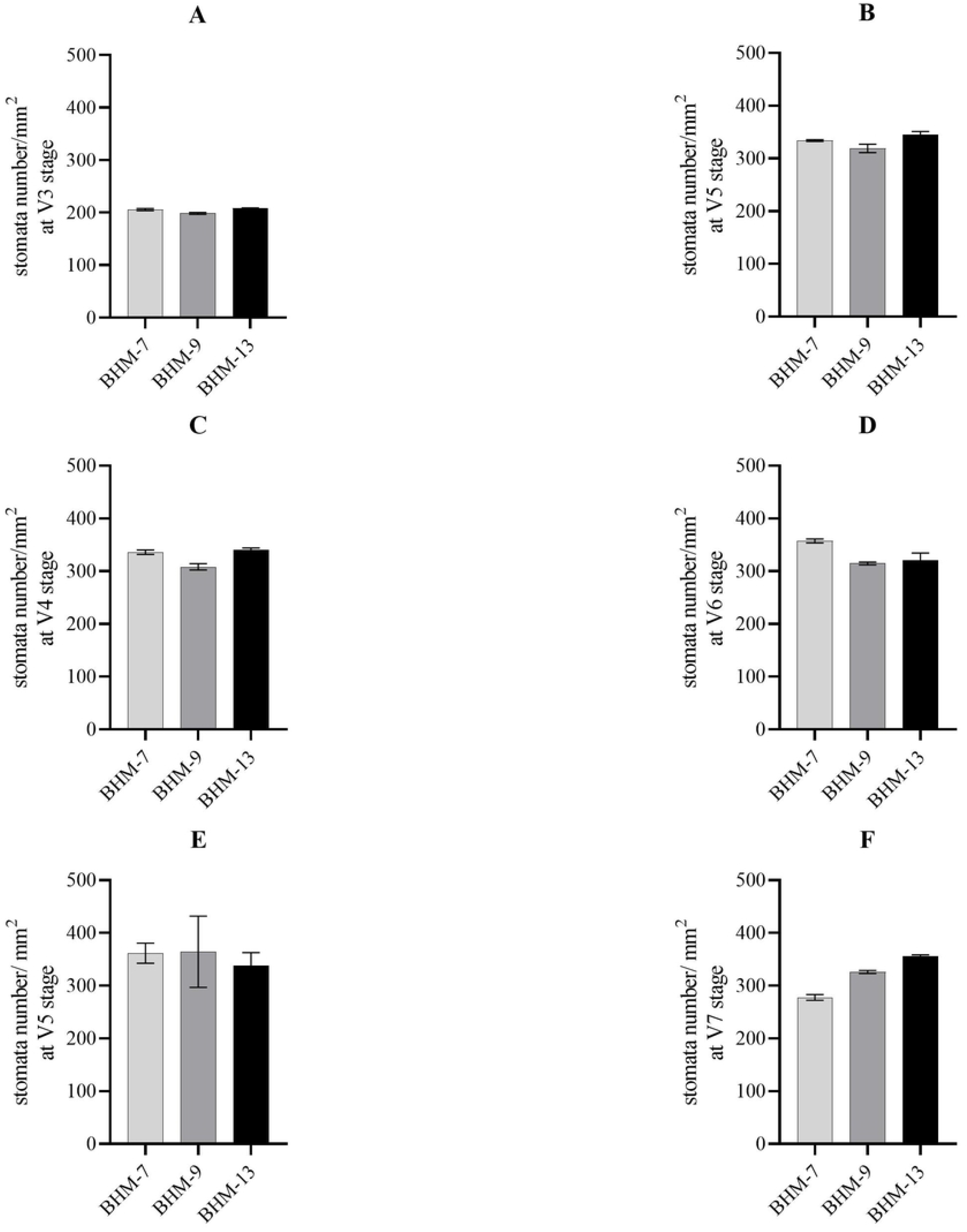
The number of stomata at two different vegetative stages of maize varieties. A. Stomatal number of BHM-7, BHM-9, BHM-13 at V3 stage before drought exposure. B. Stomatal number of BHM-7, BHM-9, BHM-13 at V5 stage before drought exposure. C. Stomatal number of BHM-7, BHM-9, BHM-13 at V4 stage after drought exposure. D. Stomatal number of BHM-7, BHM-9, BHM-13 at V6 stage after drought exposure. E. Stomatal number of BHM-7, BHM-9, BHM-13 at V5 stage after re-watering. F. Stomatal number of BHM-7, BHM-9, BHM-13 at V7 stage after re-watering. A specific area in the second leaf of each variety was randomly selected, and the number of stomata was counted in this area (S1 Fig.). Data are presented as a mean of three biological replicates, and error bars represent the standard error of the mean.

### Stomatal closure percentages among the varieties at the V4 stage after drought treatment

After drought treatment at the V4 stage, the percentage of stomatal closure of BHM-7 was 11% and 21% for control and treated plants, respectively (Fig. 2A). The rate of stomatal closure of BHM-9 was 6% and 23% for control and treated plants, respectively (Fig. 2B). Furthermore, in BHM-13, the stomatal closure percentage was 11% and 33% for control and treated plants, respectively (Fig. 2C). Therefore, the highest rate of stomatal closure was found in the BHM-13 variety compared to BHM-7 and BHM-9 (Fig. 2D).

**Fig 2.**
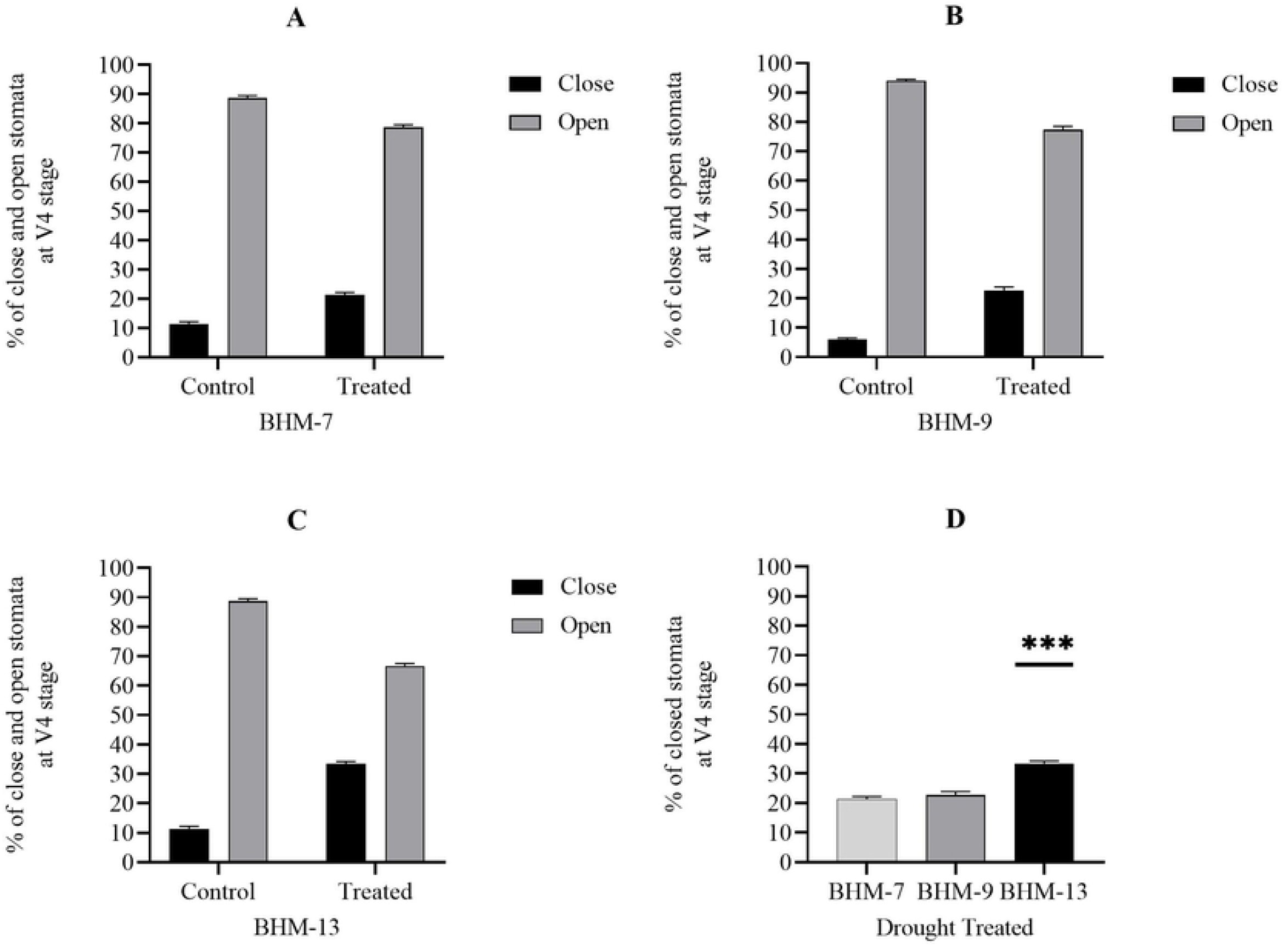
Stomatal closure and open percentage in BHM-7, BHM-9, BHM-13 at V4 stage after drought exposure. A. Stomatal closure and open percentage after drought exposure at V4 stage in BHM-7. B. Stomatal closure and open percentage after drought exposure at V4 stage in BHM-9. C. Stomatal closure and open percentage after drought exposure at V4 stage in BHM-13. D. Comparison of stomatal closure percentage of BHM-7, BHM-9, and BHM-13 at V4 stage. A specific area in the second leaf of each variety was randomly selected, and the percentage of stomatal closure was calculated in this area. Data are presented as a mean of three biological replicates, and error bars represent standard error. One-way ANOVA analysis was performed. BHM-13 showed a significantly more significant percentage of closed stomata.

### Stomatal closure percentages among the varieties at the V6 stage after drought treatment

After drought treatment at the V6 stage, the percentage of stomatal closure of BHM-7 was 14% and 18% for control and treated groups, respectively (Fig. 3A). While, for the control and treated group of BHM-9, the percentage of stomatal closure was 7% and 21%, respectively (Fig. 3B). BHM-13 had 0% and 34% closure percentages for control and treated groups, respectively (Fig. 3C). Therefore, the highest rate of stomatal closure was also found in BHM-13 compared to BHM-7 and BHM-9 at the V6 stage (Fig. 3D).

**Fig 3.**
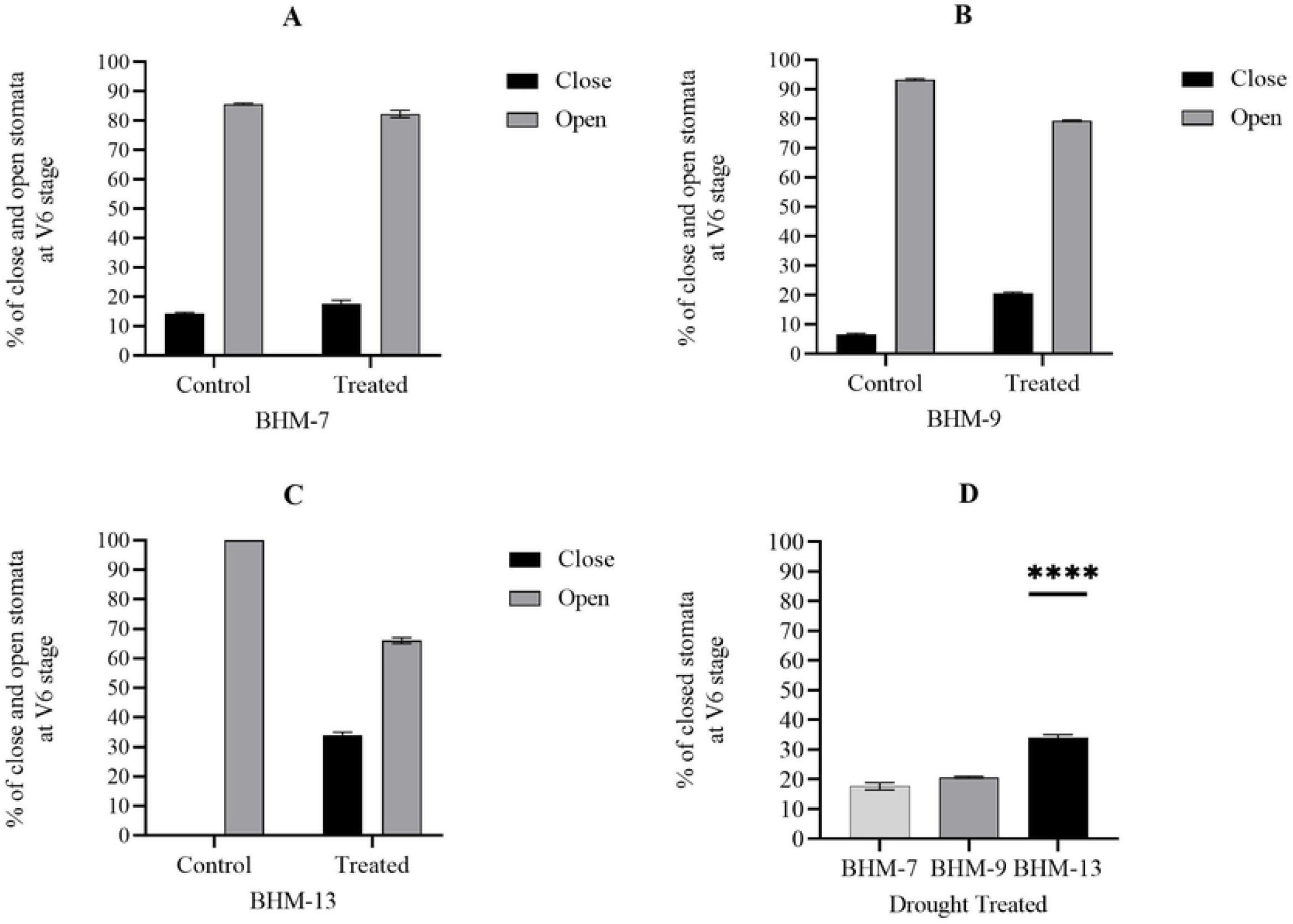
Stomatal closure and open percentage in BHM-7, BHM-9, BHM-13 at V6 stage after drought exposure. A. Stomatal closure and open percentage of BHM-7 after drought exposure at V6 stage. B. Percentage of close and open stomata after drought exposure at V6 stage in BHM-9. C. Stomatal closure and open percentage after drought treatment at V6 stage in BHM-13. D. Comparison of the percentage of stomatal closure in BHM-7, BHM-9, and BHM-13 at V6 stage. Data are presented as three biological replicates, and error bars represent the standard error. One-way ANOVA analysis was performed. BHM-13 showed a significantly more percentage of closed stomata.

### Stomatal closure percentages among the varieties at V5 and V7 stage after re-watering

After re-watering at the V5 stage, the percentages of stomatal closure of BHM-7 were 0% and 7% for control and treated groups, respectively (Fig. 4A). BHM-9 had stomatal closure percentages of 0% and 15% for control and treated plants, respectively (Fig. 4B). All of the stomata were open in BHM-13 for both control and treated groups (Fig. 4C). Moreover, after re-watering at the V7 stage, all stomata were open for all varieties for both experimental and control plants (Fig. 4D-5F).

**Fig 4.**
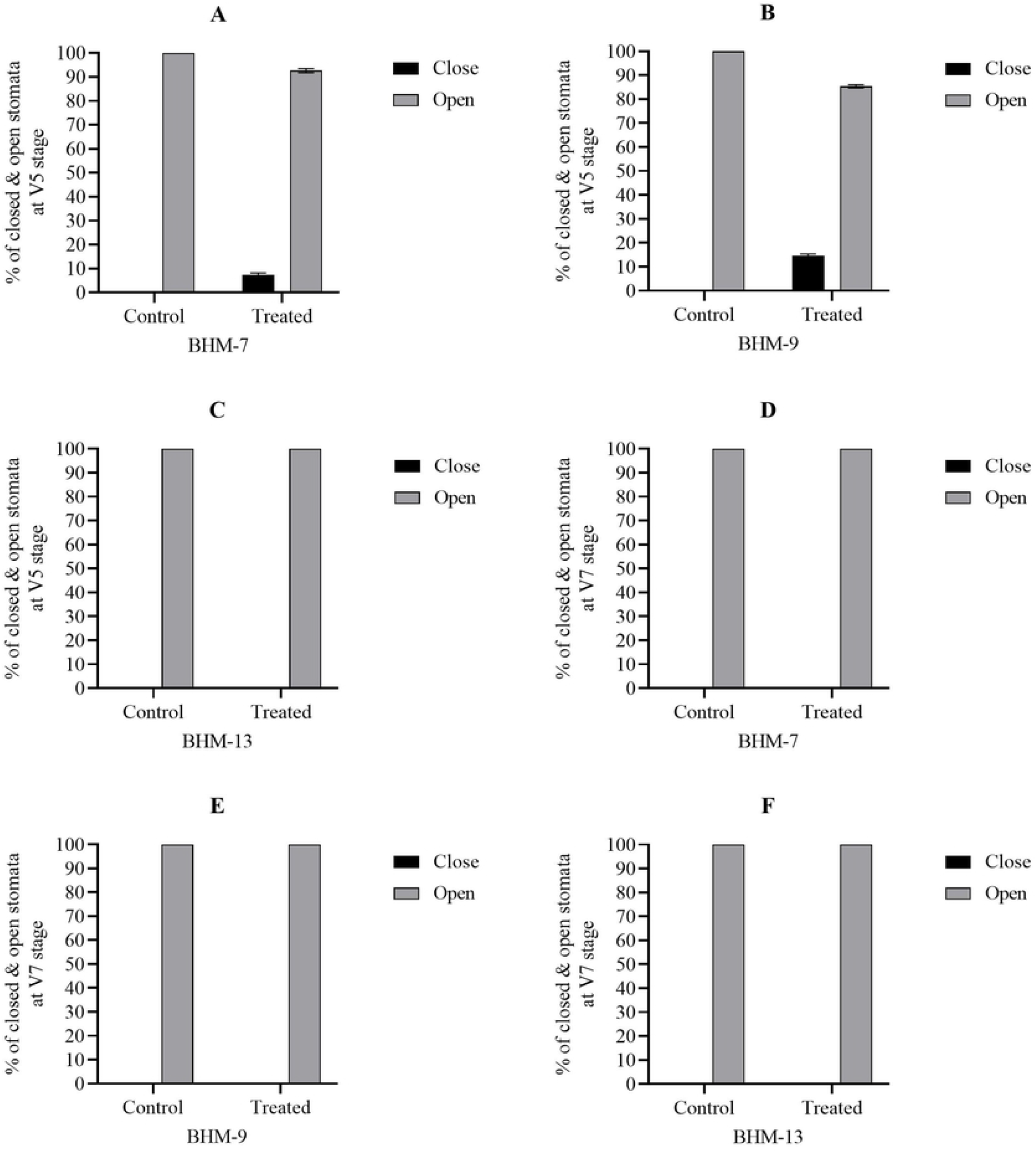
Stomatal closure and open percentage in BHM-7, BHM-9, BHM-13 at V5 and V7 stage after re-watering. A. Stomatal open & closure percentage after re-watering at V5 stage in BHM-7. B. Stomatal open & closure percentage after re-watering at V5 stage in BHM-9. C. Stomatal open & closure percentage after re-watering at V5 stage in BHM-13. D. Stomatal open & closure percentage after re-watering at V7 stage in BHM-7. E. Stomatal open & closure percentage after re-watering at V7 stage in BHM-9. F. Stomatal open & closure percentage after re-watering at V7 stage in BHM-13. Data are presented as a mean of three biological replicates, and error bars represent standard error.

### Guard cell aperture ratio among the varieties at different vegetative stages in presence or absence of drought

At V3 stage, before drought exposure the mean aperture ratio of BHM-7, BHM-9, BHM-13 was 0.051 ± 0.005 (Mean ± SE), 0.047 ± 0.005 (Mean ± SE) and 0.045 ± 0.003 (Mean ± SE), respectively (Fig. 5A). At V5 stage, prior to drought exposure the mean aperture ratio of BHM-7, BHM-9, BHM-13 was 0.073 ± 0.008 (Mean ± SE), 0.049 ± 0.005 (Mean ± SE) and 0.043 ± 0.002 (Mean ± SE), respectively (Fig. 5B).

**Fig 5.**
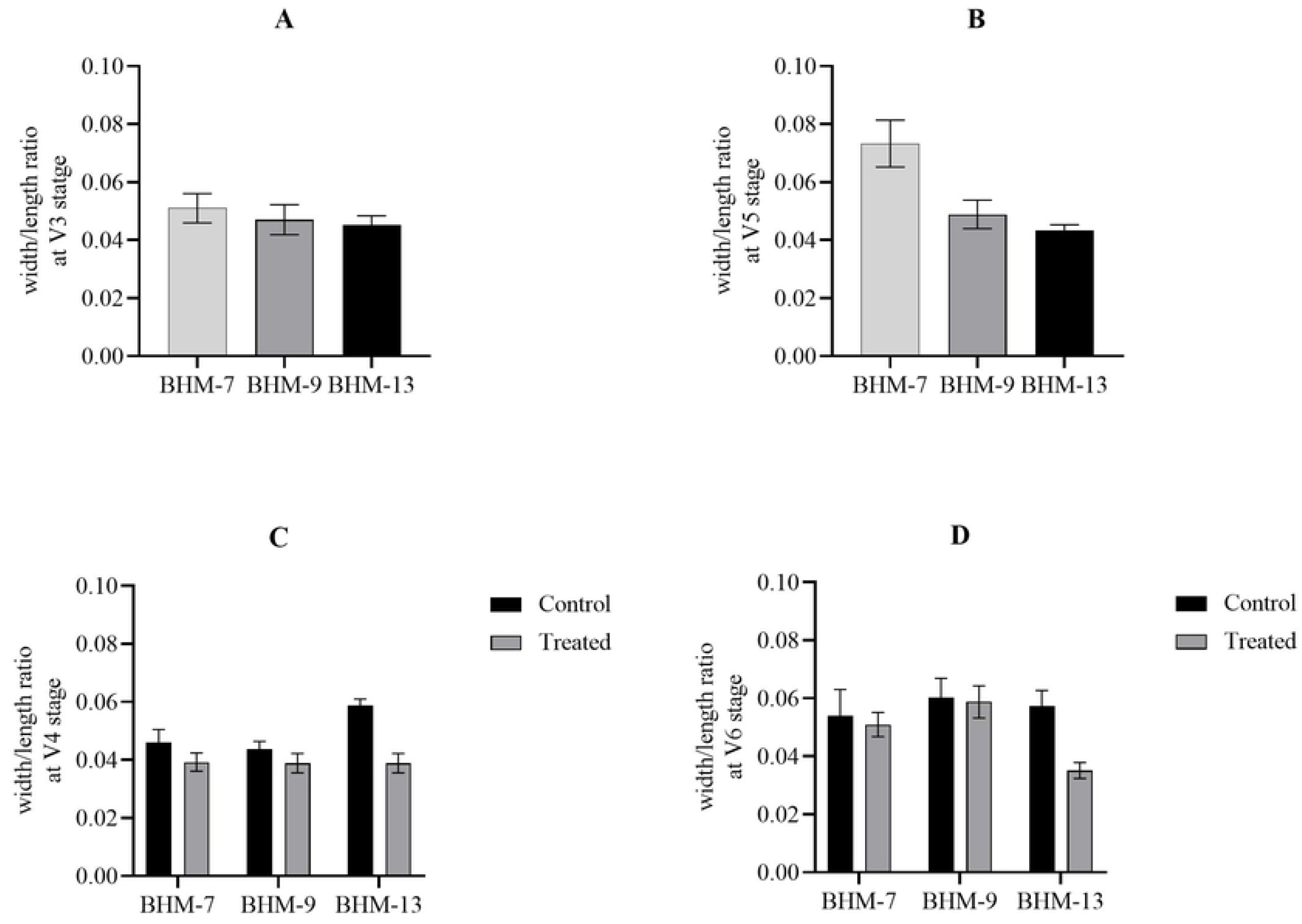
Width/Length ratio of guard cell of BHM-7, BHM-9, and BHM-13. A. Guard cell aperture ratio of BHM-7, BHM-9, and BHM-13 at V3 stage before the drought. B. Guard cell aperture ratio of BHM-7, BHM-9, and BHM-13 at V5 stage before the drought. C. Guard cell aperture ratio of BHM-7, BHM-9, and BHM-13 at V4 stage after drought treatment. D. Guard cell aperture ratio of BHM-7, BHM-9, and BHM-13 at V6 stage after drought treatment. A specific area in the second leaf of each variety was randomly selected, and guard cell aperture ratio was calculated according to Johnson R (2007) (S2 Fig.). Data are presented as a mean of three biological replicates, and error bars represent standard error.

However, after drought exposure at the V4 stage, the mean aperture ratio of BHM-7 was 0.046 ± 0.004 (Mean ± SE) and 0.039 ± 0.003 (Mean ± SE) for control and treated plants, respectively (Fig. 5C). The mean aperture ratio of BHM-9 was 0.043 ± 0.002 (Mean ± SE) and 0.038 ± 0.003 (Mean ± SE) for the control and treated plants, respectively (Fig. 5C). The mean aperture ratio of BHM-13 was 0.058 ± 0.002 (Mean ± SE) and 0.038 ± 0.003 (Mean ± SE) for the control and treated plants, respectively (Fig. 5C). The mean aperture ratio of BHM-7 and BHM-9 for control and treated plants was not significantly differ each other (p=>0.05; students’ t-test), however, a statistically significant difference was found in the BHM-13 control and treated plants (p=0.002<0.05; students’ t-test). Percentages of aperture decrease were 14.83%, 10.92%, and 33.85% in BHM-7, BHM-9, and BHM-13, respectively, upon drought treatment compared to their control groups.

At the V6 stage after 7 days of drought treatment, the mean aperture ratio of BHM-7 was 0.054 ± 0.009 (Mean ± SE), and 0.051 ± 0.004 (Mean ± SE) for control and drought treated plants, respectively (Fig. 5D). In BHM-9, the mean aperture ratio was 0.060 ± 0.007 (Mean ± SE), and 0.058 ± 0.006 (Mean ± SE) for control and drought treated plants, respectively (Fig. 5D). Whereas in BHM-13 control and drought treated plants, the mean aperture ratio was 0.057± 0.005 (Mean ± SE) and 0.035 ± 0.002 (Mean ± SE), respectively (Fig. 5D). The mean aperture ratio of BHM-7 and BHM-9 for control and treated plants was not significantly differ each other (p=>0.05; students’ t-test), however, a statistically significant difference was found in the BHM-13 control and treated plants (p=0.008<0.05; students’ t-test). The percentage of aperture reduction in BHM-7, BHM-9, and BHM-13 was 5.52%, 2.48%, and 38.75 %, respectively, upon drought treatment compared to control groups.

### Guard cell aperture ratio among the varieties after re-watering

After re-watering at the V5 stage, the mean aperture ratio of BHM-7 was 0.060 ± 0.0080 (Mean ± SE) and 0.050 ± 0.005 (Mean ± SE) for control and treated plant groups, respectively (Fig. 6A). The mean aperture ratio of BHM-9 was 0.0612 ± 0.006 (Mean ± SE) and 0.057 ± 0.006 (Mean ± SE) for the control and treated plants, respectively (Fig. 6A). The mean aperture ratio of BHM-13 was 0.056 ± 0.005 (Mean ± SE) and 0.054 ± 0.008 (Mean ± SE) for control and treated plant group, respectively (Fig. 6A). After re-watering, the differences between the aperture ratio of BHM-7 control and treated, BHM-9 control and treated, BHM-13 control and treated were 16.42%, 5.32%, and 3.27%, respectively.

**Fig 6.**
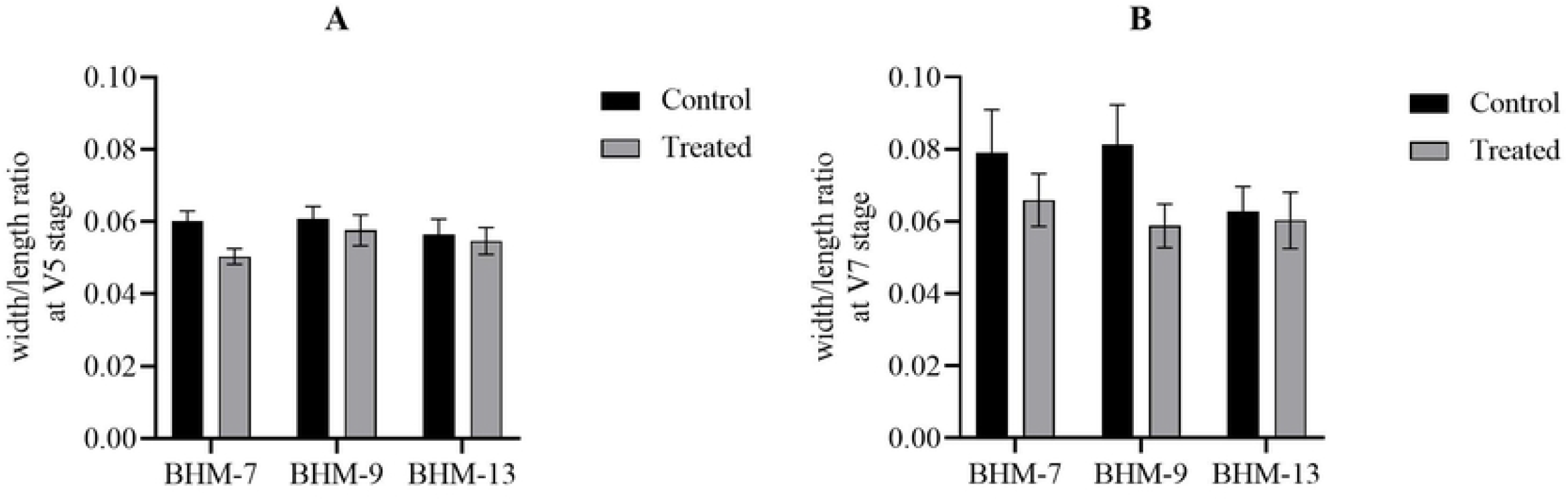
Width/Length ratio of guard cells of BHM-7, BHM-9, and BHM-13 after re-watering. A. Guard cell aperture ratio of BHM-7, BHM-9, and BHM-13 at V5 stage after re-watering. B. Guard cell aperture ratio of BHM-7, BHM-9, and BHM-13 at V7 stage after re-watering. Data are presented as a mean of three biological replicates, and error bars represent standard error.

After re-watering at the V7 stage, the mean aperture ratio of BHM-7 was 0.078±0.0079 (Mean ± SE) and 0.065±0.007 (Mean ± SE) for the control and treated groups, respectively (Fig. 6B). The mean aperture ratio of BHM-9 was 0.0612±0.006 (Mean ± SE) and 0.058±0.006 (Mean ± SE) for the control and treated groups, respectively (Fig. 6B). The mean aperture ratio of BHM-13 was 0.0561±0.0058 (Mean ± SE) and 0.0550±0.0078 (Mean ± SE) for the control and treated groups, respectively (Fig. 6B). After re-watering, the differences between aperture ratio of BHM-7 control and treated, BHM-9 control and treated, BHM-13 control and treated were 16.50%, 27.74%, and 4.03%, respectively. Both at V5 and V7 stage, BHM-13 showed the lowest percentages of aperture ratio compared to BHM-7 and BHM-9 indicating that BHM-13 has high water use efficiency compared to other varieties.

## Discussion

### The stomatal number varies at different vegetative stages

The stomatal density of BHM-7, BHM-9 and BHM-13 varies each other with the age of plants and treatment condition (Fig. 1). Stomatal density was increased in all plants in V4 and V5 stage (Fig. 1C and Fig. 1E) compared to V3 (Fig. 1A). However, stomatal density was decreased in BHM-7 at V7 stage (Fig. 1F) compared to V5 (Fig. 1B) and V6 stage (Fig. 1D). Drought has a strong influence in stomatal density, guard cell size and aperture ratio. It has been found that severe drought can lead to a reduction in stomatal number, though an increase in stomatal number is possible under low or moderate drought conditions (16). The responses of guard cell size and stomatal number to environmental variables depend on a time scale from milliseconds to millions of years (17). The physiological mechanisms of stomatal response are very complex and not yet fully understood to date (18, 19).

### BHM-13 showed the highest stomatal closure and lowest aperture ratio in both vegetative stages under drought exposure

After 7 days’ drought treatment, BHM-13 showed highest percentage of closed stomata compared to BHM-7 and BHM-9 both at the V4 stage (Fig. 2C) and at the V6 stage (Fig. 3C). By synthesizing abscisic acid, it effectively reduces water loss in drought conditions than the other two varieties regarding stomata closing phenomena (20). After re-watering for 7 days, no stomata were closed in BHM-13 (Fig. 4C), however, BHM-7 and BHM-9 had 7% and 15% closed stomata, respectively at V5 stage (Fig. 4A and Fig. 4B). This was due to a physiologically younger form of leaves of stressed plants following turgor regaining (21). During endosmosis or the entry of water, stomata became turgid, which resulted in stomatal opening. When the turgor develops within the two guard cells, the thin outer walls bulge outward and force the inner walls into a crescent shape to open the stomata. This is the state during which the exchange of oxygen, carbon dioxide, and water vapor loss occurred through pores (3, 22).

At the V3 stage and V5 stage, before the drought, BHM-13 had the lowest aperture ratio (Fig. 5A and 5B). After drought exposure at V4 and V6 stage, a highest reduction of the aperture ratio was found in BHM-13 treated plants compared to control plants (Fig. 5C and 5D). This is probably due to the production of high abscisic acid in the treated plants under water deficit condition, that subsequently leads to a reduction of the aperture between guard cells (20). A similar aperture ration was found in control and treated plants, both for BHM-7 and BHM-9 (Fig. 5C and 5D). After drought exposure followed by re-watering, the aperture ratio of the treated group was very close to the control group for BHM-13 (Fig. 6A and 6B). However, the aperture ratio is varied for the control and treated plants, both for the BHM-7 and BHM-9 (Fig. 6A and 6B). Plants’ abiotic stress adaptation mechanisms are very complex; however, abscisic acid is the crucial regulator of adaptation mechanisms (23).

Therefore, BHM-13 is the most drought-resistant variety amongst the three tested varieties (BHM-7, BHM-9, and BHM-13). It has been found that BHM-13 is morphologically short and capable of developing side branching from its lower nodes under heat stress condition (24). Since drought-resistant plants should combine a better root system, stomatal regulation, water-use efficiency, and hormonal balance, further morphological, biochemical & yield analyses are recommended to find out the drought-resistant variety more precisely.

## Acknowledgments

We thank Biotechnology and Genetic Engineering Discipline and Agrotechnology Discipline, Khulna University for their institutional and technical support.

## Author Contributions

Conceived and designed the experiment: SMAA. Performed the experiment: MMUT, PD. Analyzed the data: SN, MRA. Contributed to instrumental and analytical tools: MRA, MRI. Wrote the original draft of Manuscript: MMUT, SA, SN. Review and editing of the draft of the Manuscript: SMAA.

## Supporting Information

**S1 Fig. Stomatal counting of leaf epidermal peel.**

**S2 Fig. Measurement of guard cell aperture ratio.**

